# Treatment of mice with IL2-complex enhances inflammasome-driven IFN-γ production and prevents lethal toxoplasmosis

**DOI:** 10.1101/593574

**Authors:** Andreas Kupz, Saparna Pai, Paul R. Giacomin, Jennifer A. Whan, Robert A. Walker, Pierre-Mehdi Hammoudi, Nicholas C. Smith, Catherine M. Miller

## Abstract

Toxoplasmic encephalitis is an AIDS-defining condition in HIV^+^ individuals. The decline of IFN-γ-producing CD4^+^ T cells in AIDS is a major contributing factor in reactivation of quiescent *Toxoplasma gondii* to an actively replicating stage of infection. Hence, it is important to identify CD4-independent mechanisms to control acute *T. gondii* infection. Here we have investigated the targeted expansion and regulation of IFN-γ production by CD8^+^ T cells, DN T cells and NK cells in response to *T. gondii* infection using IL-2 complex (IL2C) pre-treatment in an acute *in vivo* mouse model. Our results show that expansion of CD8^+^ T cells, DN T cells and NK cell by S4B6 IL2C treatment increases survival rates of mice infected with *T. gondii* and this increased survival is dependent on both IL-12- and IL-18-driven IFN-γ production. Processing and secretion of IFN-γ-inducing, bioactive IL-18 is dependent on the sensing of active parasite invasion by multiple redundant inflammasome sensors in multiple hematopoietic cell types but independent from *T. gondii*-derived dense granule (GRA) proteins. Our results provide evidence for a protective role of IL2C-mediated expansion of CD8^+^ T cells, DN T cells and NK cells in murine toxoplasmosis and may represent a promising adjunct therapy for acute toxoplasmosis.

**Author Summary:** A third of the world’s population is chronically infected with the parasite *Toxoplasma gondii*. In most cases the infection is asymptomatic, but in individuals suffering from AIDS, reactivation of brain and muscle cysts containing *T. gondii* is a significant cause of death. The gradual decline of CD4 T cells, the hallmark of AIDS, is believed to be a major contributing factor in reactivation of *T. gondii* infection and the development of acute disease. In this study, we show that targeted expansion of non-CD4 immune cell subsets can prevent severe disease and premature death via increased availability of interferon gamma-producing immune cells. We also demonstrate that the upstream signaling molecule interleukin-18 is required for the protective immune response by non-CD4 cells and show that the sensing of active parasite invasion by danger recognition molecules is crucial. Our findings reveal that targeted cell expansion may be a promising therapy in toxoplasmosis and suggests that the development of novel intervention strategies targeting danger recognition pathways may be useful against toxoplasmosis, particularly in the context of AIDS.

## Introduction

*Toxoplasma gondii* (*T. gondii*) is an obligate intracellular parasite of the phylum Apicomplexa [1]. It is estimated that one-third of the world’s population is infected with *T. gondii*. In most individuals, infection is asymptomatic and leads to chronic, life-long persistence of *T. gondii*-containing cysts, primarily in brain and muscle tissue [2]. Active disease, also known as toxoplasmosis, usually occurs after reactivation of encysted parasites, and is often associated with immunosuppression. If untreated, toxoplasmosis may be fatal. Additionally, serious eye disease has been reported as a result of infection with *T. gondii* [3] and, if a primary infection occurs during pregnancy, abortion, stillbirth and fetal abnormalities can occur [2, 4]. Whereas an acute infection is generally mediated by the fast-replicating tachyzoite stage of the parasite, the persistent tissue cysts, characteristic of a chronic infection, contain slow-replicating bradyzoites. Currently, treatment of toxoplasmosis is limited to the acute disease and requires prolonged exposure to anti-toxoplasmosis drugs for the duration of the immunosuppression [5, 6].

Containment of chronic *T. gondii* infection requires functional T-cell responses, in particular interferon gamma (IFN-γ)-producing CD4^+^ T cells [2, 7]. In the absence of CD4^+^ T cells, IFN-γ, its receptor or downstream effector molecules, such as inducible nitric oxide synthase (iNOS), susceptibility and disease are severely exacerbated [8–11]. Accordingly, co-infection with human immunodeficiency virus (HIV), which impairs CD4^+^ T cells during its reproduction, is one of the major reactivation factors. In fact, toxoplasmic encephalitis accompanied by low numbers of CD4^+^ T cells is considered to be an AIDS-defining condition in HIV^+^ individuals [12].

In addition to antigen-specific CD4^+^ T cells [11], innate immune cells, such as NK cells and neutrophils also contribute significantly to the production of host-protective IFN-γ [13, 14]. In particular, the recognition of *T. gondii*-derived profilin via Toll-like receptor (TLR)-11, which drives myeloid differentiation primary-response protein 88 (MyD88)-dependent IL-12 secretion by dendritic cells, is considered a crucial upstream pathway of protective IFN-γ secretion [15, 16]. Mice deficient in MyD88 or IL-12 are also extremely susceptible to *T. gondii* infection [17, 18]. Furthermore, elegant studies by Hunter and colleagues showed that T cell-intrinsic ablation of MyD88 also impacts severely on the control of the parasite [19]. These findings indicate that, in addition to IL-12, cytokine-driven IFN-γ secretion in response to *T. gondii* also depends on IL-18, an IL-1 family cytokine originally known as IFN-γ-inducing factor, which requires cell-intrinsic MyD88 signaling [20, 21]. IL-18 is particularly important for the rapid production of IFN-γ by cells of the immune system, in particular NK cells, CD8^+^ memory T cells and double negative (DN) γδ T cells [22].

Secretion of bioactive IL-18 requires proteolytic cleavage from its biologically inactive precursor, pro-IL-18, through caspase-1 [23], which in turn depends on the upstream assembly and activation of inflammasomes through the engagement of cytosolic pattern recognition receptors (PRRs) [23]. Intriguingly, not only deficiencies in caspase-1 and IL-18 [24, 25] have been implicated in impaired immunity to *T. gondii*, but also deficiencies in the inflammasome sensors NLRP1 and NLRP3 [24, 26]. These results point to an important host-protective role for the caspase1 → IL-18 → IFN-γ axis and suggest that strategies aimed at targeting cytosolic PRRs as adjunct immunotherapy [27] could serve as a means of inducing IL-18-mediated IFN-γ production to control infections with *T. gondii*. Consistent with this hypothesis, we and others have recently demonstrated, in models of experimental *Listeria monocytogenes, Mycobacterium tuberculosis* and *Salmonella enterica* infection, that rapid, IL-18-driven IFN-γ secretion orchestrates host innate immunity and impacts on the magnitude of the recall response after vaccination [28–30].

Given that control of acute toxoplasmosis depends on a delicate balance between limiting immunopathology and maintaining parasite killing, in the present study, we interrogated the mechanistic regulation of IL-18-driven IFN-γ production *in vivo*. We discovered that bioactive IL-18 is dependent on the sensing of active parasite invasion by multiple redundant inflammasome sensors in multiple non-CD4 hematopoietic cell types, leading to the hypothesis that enhancement of this innate response could be harnessed to prevent disease resulting from infection with *T. gondii*. We therefore investigated if treatment with S4B6-containing IL2C, an IL2 complex that can boost NK and CD8^+^ T cell numbers [31], could prevent acute lethal toxoplasmosis.

## RESULTS

### *Toxoplasma*-driven IFN-γ secretion by non-CD4 immune cells following oral infection with brain cysts or intravenous (i.v.) infection with tachyzoites

Given that control of acute toxoplasmosis critically depends on IFN-γ [7] and non-CD4 immune cell types, such as CD8^+^ T cells, DN T cells and NK cells, are prime IFN-γ producers, we wanted to delineate the mechanistic requirements of IFN-γ production by these cell types in response to *T. gondii*. We furthermore wanted to explore whether responses were similar after oral infection (a common natural route of infection), i.v. infection with tachyzoites (modelling blood transfusion, a rare but significant – for the individual – route of infection [32]) and the often used purely experimental i.p. route of infection with tachyzoites.

We first inoculated naïve B6 mice with 10, 40 or 100 *T. gondii* ME49 cysts and assessed IFN-γ production by viable splenic CD3^+^CD4^+^, CD3^+^CD8^+^, CD3^+^CD4^−^CD8^−^ (DN) T cells and CD3^−^ NKp46^+^ cells 1 day and 5 days after inoculation. Whereas no IFN-γ production was observed 1 day after inoculation, a significant increase in IFN-γ-secreting cells was detected at 5 days after inoculation in spleen, MLN and PP (**Fig 1A, B and S1A, B Fig**). Up to 10% of CD8^+^ T cells and DN T Cells and up to 50% of all NK cells stained IFN-γ^+^, particularly following inoculation with 40 and 100 cysts. Because these mice had never been exposed to apicomplexan parasites before, these results ruled out antigen-specific responses.

**Figure 1:**
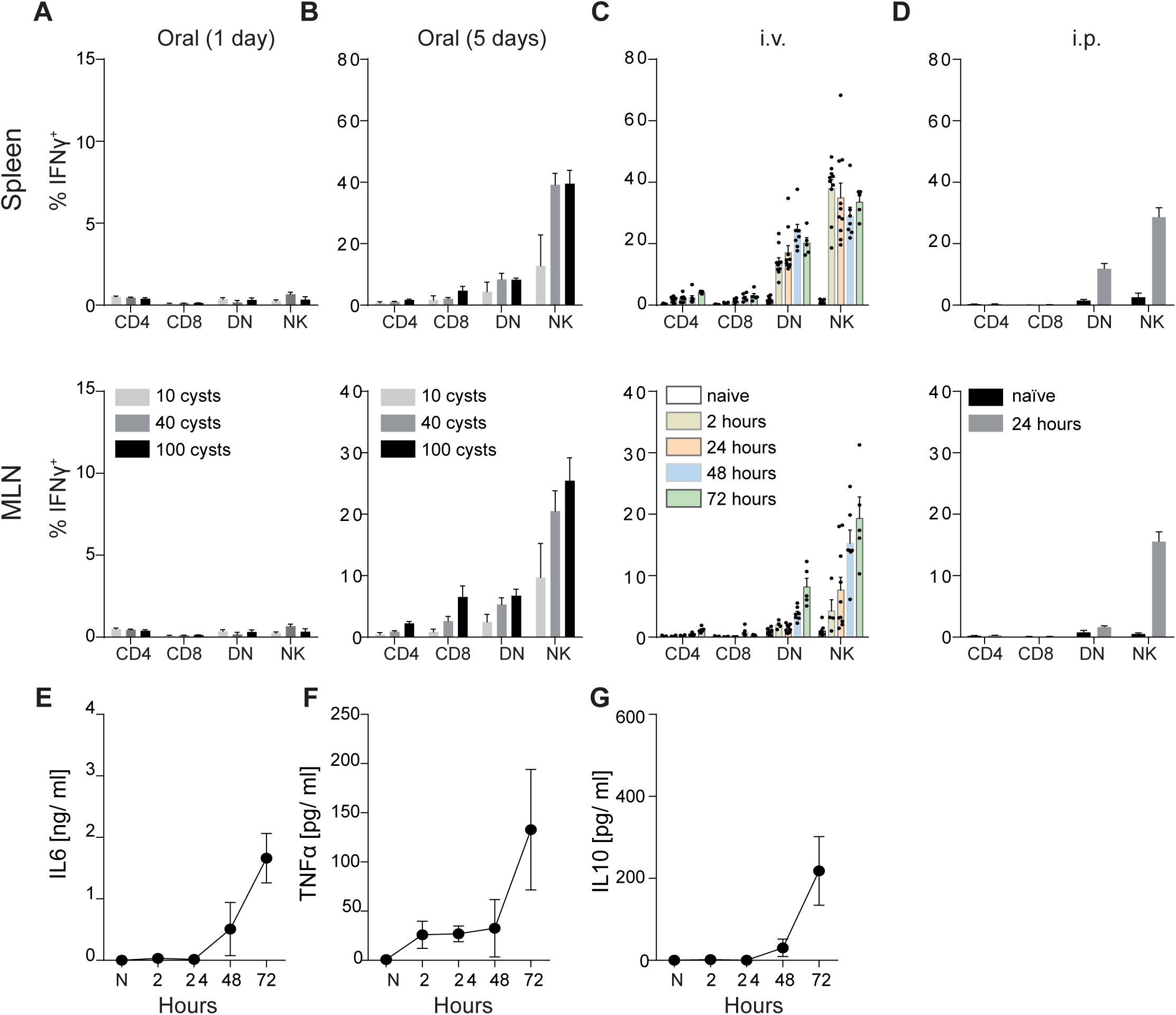
*Toxoplasma*-driven IFN-γ secretion by non-CD4 immune cells following oral infection with brain cysts or intravenous (i.v.) infection with tachyzoites. (A, B) Percent of IFN-γ^+^ cells amongst total viable splenic CD3^+^CD4^+^, CD3^+^CD8^+^, CD3^+^CD4^−^CD8^−^ (DN) T cells and CD3^−^NKp46^+^ cells 1 day (A) or 5 days (B) after B6 mice were inoculated orally with 10, 40 or 100 *T. gondii* ME49 brain cysts. (C) Percent of IFN-γ^+^ cells amongst total viable splenic CD3^+^CD4^+^, CD3^+^CD8^+^, CD3^+^CD4^−^CD8^−^ (DN) T cells and CD3^−^NKp46^+^ cells at 2-72 hours after B6 mice were injected i.v. with 10^7^ *T. gondii* ME49 tachyzoites. (D) Percent of IFN-γ^+^ cells amongst total viable splenic CD3^+^CD4^+^, CD3^+^CD8^+^, CD3^+^CD4^−^CD8^−^ (DN) T cells and CD3^−^NKp46^+^ cells at 24 hours after B6 mice were injected i.p. with 10^7^ *T. gondii* ME49 tachyzoites. (E-G) Serum concentrations of IL-6 (E), TNFα (F) and IL-10 (G) at 2-72 hours after B6 mice were injected i.v. with 10^7^ *T. gondii* ME49 tachyzoites. Results are presented as pooled data means ± SEM from at least two pooled independent experiments (n = 5-10 mice per group). See also S1 Figure.

We also investigated if rapid IFN-γ production could be induced by inoculation with tachyzoites via the i.v. and i.p. routes using a short-term *in vivo* exposure model in which naïve B6 mice were exposed to *T. gondii* tachyzoites for a maximum of 72 hours. When mice were injected i.v. or i.p. with 10^5^ tachyzoites, no significant IFN-γ production could be seen in either spleen, MLN or PP within 72 hours (**S1E Fig**). However, i.v. or i.p. inoculation with 10^7^ tachyzoites led to secretion of IFN-γ by CD3^+^CD8^+^, CD3^+^CD4^−^CD8^−^ (DN) T cells and CD3^−^NKp46^+^ cells in spleen, MLN and PP as early as 2-24 hours after inoculation (**Fig 1C, D and S1C, D Fig**), mirroring the results seen 5 days after a cyst inoculation (**Fig 1B**). Importantly, at 24 hours after tachyzoite inoculation, levels of other acute inflammatory mediators, such as IL-6, TNFα and IL-10, were almost indistinguishable from naïve mice (**Fig 1E-G**). These results indicate that mice were still controlling the infection and that parasite dissemination and subsequent acute cytokine responses were not yet impacting on protective IFN-γ responses 24 hours after i.v. infection.

Furthermore, these results show that i.v., i.p. tachyzoite infections and oral brain cyst infections induce almost identical acute immune responses. *Toxoplasma gondii* cyst production in mice is a slow and laborious process. In addition, it is difficult to quantify the number of bradyzoites within brain cysts used for oral infection and, moreover, dissemination patterns following oral infection are erratic in individual mice [33]. Therefore, we subsequently focused on IFN-γ secretion by splenic NK cells 24 hours after i.v. injection of tachyzoites as our primary readout for further dissection of the underlying mechanistic requirements.

### Rapid IFN-γ secretion in response to *T. gondii* requires IL-12 & IL-18

Whereas the role of IL-12 in IFN-γ secretion is well established for *T. gondii* [2], rapid production of IFN-γ in response to other intracellular pathogens, such as *S. enterica, L. monocytogenes* and *M. tuberculosis* has also been linked to the upstream effects of IL-18 [28, 29]. To interrogate whether or not, and how early, IFN-γ secretion in response to *T. gondii* also requires IL-18, we exposed naïve B6 mice to *T. gondii* ME49 tachyzoites and treated the animals with neutralizing monoclonal antibodies (mAb) to IL-12, IL-18 or IL-12 and IL-18 immediately after inoculation. At 24 hours after exposure, IFN-γ secretion by NK cells in the spleen was assessed directly *ex vivo*. Neutralization of IL-12 and IL-18 significantly reduced IFN-γ production, with IL-12 contributing approximately 50% and IL-18 approximately 30-40% of the response (**Fig 2A**). The significant reduction of rapid IFN-γ production in *Il18*^*–/–*^ mice, and the almost complete absence of rapid IFN-γ production in anti-IL-12-treated *Il18*^*–/–*^ mice, further confirmed a direct correlation between IL-12, IL-18 and IFN-γ secretion (**Fig 2C, D**). Consistently, where IL-12 levels in the serum of infected mice peaked at approximately 2 hours after inoculation, the levels of IL-18 mirrored those of IFN-γ for up to 72 hours (**Fig 2B**). Furthermore, treatment with anti-IL-12 and/or anti-IL-18 also reduced concentrations of IFN-γ, IL-12 and IL-18 in the serum of infected mice in an additive manner (**Fig 2D-F**). These results suggest a hierarchical relationship in which a primary IL-12-driven IFN-γ response is followed by an IL-18-dominant IFN-γ response. We concluded that innate IFN-γ secretion by CD8^+^ T cells, DN T cells and NK cells in response to *T. gondii* is driven by the secretion of IL-12 and IL-18.

**Figure 2:**
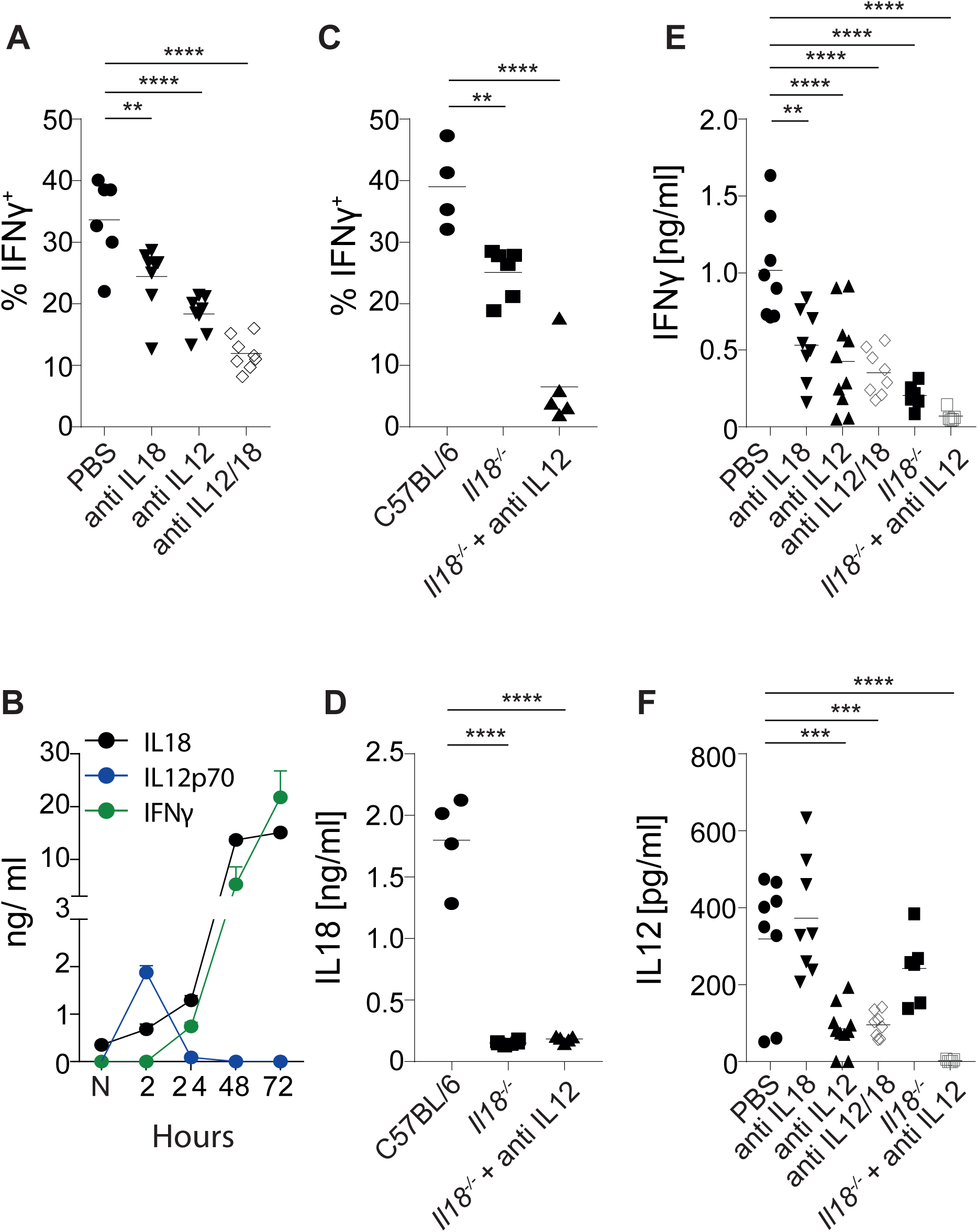
Rapid IFN-γ production in response to *T. gondii* requires IL-12 and IL-18. Percent of IFN-γ^+^ cells amongst total viable CD3^−^NKp46^+^ cells in the spleen (A, C) and serum cytokine concentrations (D-F) 24 hours after B6 or *Il18*^*-/*-^ mice were injected i.v. with 10^7^ *T. gondii* ME49 tachyzoites. Some mice received an i.p. injection of 200 µg mAb against IL-18 and/or IL-12 immediately after injection of *T. gondii*. (B) Serum concentrations of IL-18, IL-12p70 and IFN-γ at various time points after i.v. injection of 10^7^ *T. gondii* ME49 tachyzoites. Some mice were additionally treated with mAb against IL-12 and/or IL-18 immediately after injection of *T. gondii* ME49. Results are presented as individual data points (A, C, D-F) or as means ± SEM (B) of 4-15 mice per group from at least two pooled independent experiments. Statistical analyses: One-way ANOVA followed by Dunnett’s multiple comparison test; significant differences are indicated by asterisks: * p<0.05; ** p<0.01; *** p<0.001; **** p<0.0001.

### IL-18-driven IFN-γ secretion to *T. gondii* depends on multiple redundant inflammasomes

Given that the molecular mechanisms that lead to *T. gondii*-mediated IL-12 secretion are well characterized, we focused our attention on the host signaling pathways required for IL-18-driven IFN-γ production, using a panel of genetically modified mouse strains. Secretion of bioactive IL-18 depends on the enzymatic cleavage of pro-IL-18 by caspase-1 [23]. Activation of caspase-1 involves the sensing of danger molecules or stress signals *via* upstream cytosolic PRRs, so called inflammasomes, a process that can be enhanced and controlled *via* TRIF-dependent caspase-11 activation. *Caspase1/11*^*–/–*^ double KO mice produced significantly less IFN-γ following injection with *T. gondii* ME49 tachyzoites compared with B6 mice, and this response could be almost completely prevented by additional anti-IL-12 treatment (**Fig 3A**). As expected, *Caspase1/11*^*–/–*^ mice did not secrete significant levels of IL-18 following *T. gondii* inoculation (**Fig 3B**), indicating that the remaining IFN-γ response in *Caspase1/11*^*–/–*^ mice is driven by IL-12. Surprisingly, when we tested mice deficient in the upstream NLR family pyrin domain-containing proteins 1 and 3 (NLRP1 and NLRP3), NLR molecules that had been implicated previously in recognition of *T. gondii* [24], both knockout strains secreted indistinguishable amounts of IL-18 compared with B6 mice (**Fig 3B**). This data suggested a redundant role for NLRP1 and NLRP3. However, even double knockout and heterozygous *Nlrp1*^*±/-*^*Nlrp3*^*±/-*^ mice secreted high levels of IL-18 and IFN-γ after exposure to *T. gondii* ME49 tachyzoites (**Fig 3A, B**), suggesting that additional PRR molecules must be involved in sensing of *T. gondii* invasion *in vivo*. Taken together these results indicate that rapid IFN-γ secretion *in vivo* in response to *T. gondii* depends on the inflammasome → caspase-1 → IL-18 axis, and that *T. gondii* activates at least three different inflammasomes *in vivo*.

**Figure 3:**
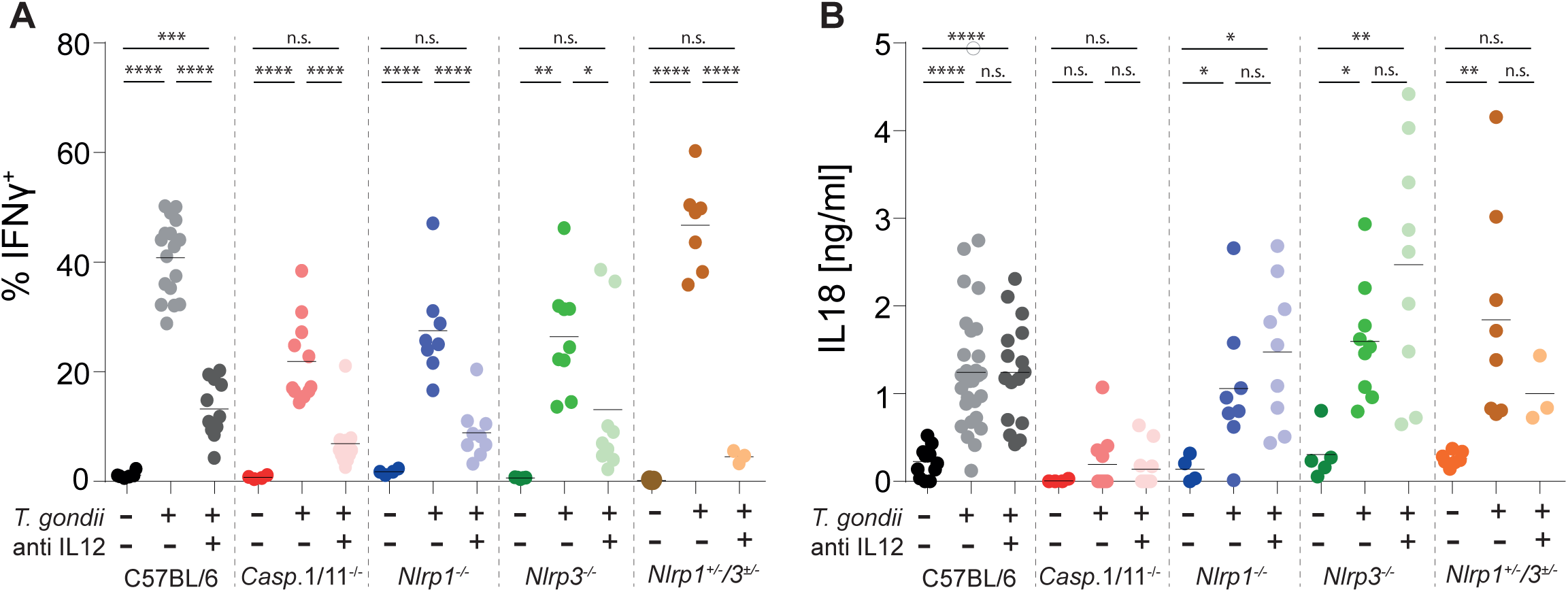
IL-18-driven IFN-γ secretion to *T. gondii* depends on multiple redundant inflammasomes. (A, B) Percent of IFN-γ^+^ cells amongst total CD3^−^NKp46^+^ cells in the spleen (A) and serum IL-18 concentrations (B) 24 hours after i.v. injection of 10^7^ *T. gondii* ME49 tachyzoites into B6 mice and different mouse strains lacking either *Caspase1/11, Nlrp1, Nlrp3* or *Nlrp1* and *Nlrp3*. Results are presented as individual data points of 3-25 mice per group from at least two pooled independent experiments. Statistical analyses: One-way ANOVA per strain followed by Dunnett’s multiple comparison test; significant differences are indicated by asterisks: * p<0.05; ** p<0.01; *** p<0.001; **** p<0.0001; n.s. not significant.

### *Toxoplasma gondii* activates inflammasomes in multiple cell types

To further investigate the role of cytosolic PRRs in sensing *T. gondii* invasion, and to potentially target inflammasome activation for preventive or therapeutic intervention strategies, we next tried to identify the *T. gondii*-sensing cell type *in vivo*. To do this, we made use of a red fluorescent protein (RFP) tagged *T. gondii* ME49 (*T. gondii* ME49-RFP) strain to track parasite uptake by different immune cell subsets in the spleen. Twenty-four hours after tachyzoite injection, *T. gondii* ME49-RFP also induced rapid IFN-γ secretion by splenic CD3^+^CD4^+^, CD3^+^CD8^+^, CD3^+^CD4^−^CD8^−^ (DN) T cells and CD3^−^NKp46^+^ cells (**Fig 4A**) and high levels of serum IL-18 (**Fig 4B**), similar to wild-type *T. gondii* ME49 (see Figs. 1 and 2). Approximately 0.5% of all splenocytes contained *T. gondii* ME49-RFP *in vivo* 24 hours after inoculation (**Fig 4C**). Sorted RFP^+^ cells secreted significantly more IL-18 *ex vivo* compared to RFP^-^ cells (**Fig 4D**), and further surface phenotyping revealed that *T. gondii* ME49-RFP was primarily contained in monocytes, neutrophils and CD8α^+^ dendritic cells (**Fig 4E, F**). Splenic MHC-II^+^CD11c^+^ DCs, CD11b^+^Ly6G^+^ neutrophils and CD11b^+^Ly6C^+^ monocytes each comprised approximately 20-30% of all RFP-containing cells after i.v. tachyzoite injection. Only very few T cells, B cells and macrophages appeared to harbor parasites (**Fig 4E, F**). To investigate if cell types that contained *T. gondii* ME49-RFP parasites also activated inflammasomes, we performed intracellular staining for the inflammasome adaptor molecule apoptosis-associated speck-like protein containing a carboxy-terminal CARD (ASC), and measured the activation of caspase-1 with a fluorescent inhibitor that only binds to activated caspase-1 (FLICA FAM-YVAD-FMK) [29]. Consistent with the uptake of *T. gondii* ME49-RFP by different cell types, *T. gondii* ME49-RFP parasite-harboring neutrophils, monocytes and DCs also expressed higher levels of ASC and FAM-YVAD compared with RFP^-^ cells and FMO controls (**Fig 4G**). Collectively, these results indicate that *T. gondii* infection activates multiple redundant inflammasomes in multiple different hematopoietic cell-types *in vivo*.

**Figure 4:**
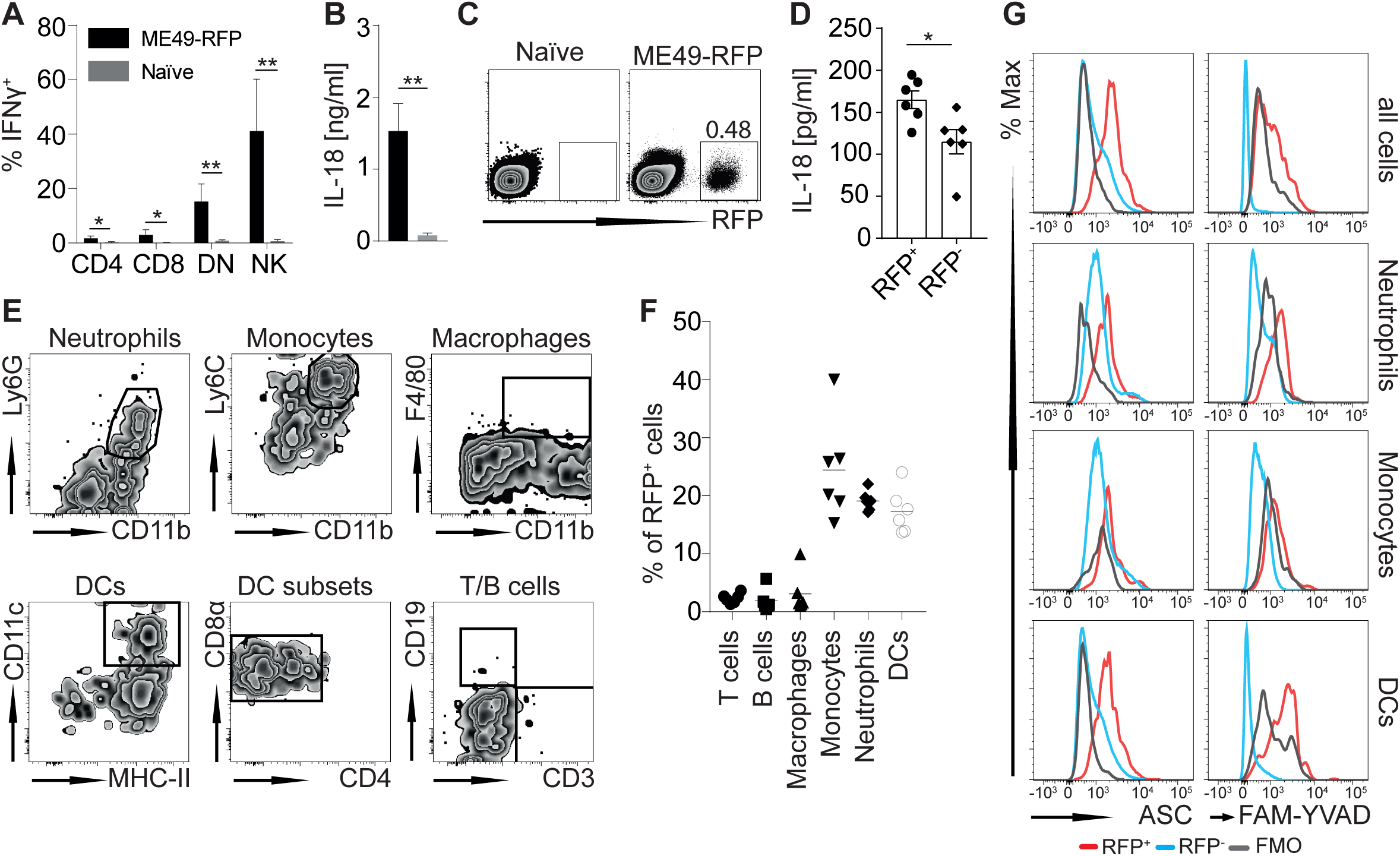
*T. gondii* activates inflammasomes in multiple cell types. (A, B) Percent of IFN-γ^+^ cells amongst total viable splenic CD3^+^CD4^+^, CD3^+^CD8^+^, CD3^+^CD4^−^CD8^−^ (DN) T cells and CD3^−^NKp46^+^ cells (A) and serum IL-18 levels (B) in naïve mice 24 hours after i.v. injection of 10^7^ *T. gondii* ME49-RFP tachyzoites. (C, E) Representative FACS plots showing total viable splenic RFP^+^ cells (C) and gated RFP^+^ cells (E) 24 hours after i.v. injection of 10^7^ *T. gondii* ME49-RFP tachyzoites. (D) IL-18 levels in supernatant of sorted RFP^+^ and RFP^-^ cells after incubation at 37°C for 24 hours. (F) Enumeration of RFP^+^ cell types shown in d. (G) Representative histograms of cell type-specific gated RFP^+^ and RFP^-^ cells showing expression levels of ASC (left panels) or FAM-YVAD (right panels) 24 hours after i.v. injection with 10^7^ *T. gondii* ME49-RFP tachyzoites. FMO control for ASC panels are cells from infected animals that did not get stained with anti-ASC-Alexa488 but all other antibodies. FMO control for FAM-YVAD are cells from mice that were injected with *T. gondii* ME49-RFP but did not receive an injection with FLICA FAM-YVAD. Results are presented as individual data points (D, F), pooled data means ± SEM (A, B) and representative FACS plots (C, E) and histograms (G) of 6-9 mice from two or three pooled independent experiments. Statistical analyses: One-way ANOVA followed by Dunnett’s multiple comparison test (A) or Student’s *t*-test (B, D); significant differences are indicated by asterisks: * p<0.05; ** p<0.01.

### IL-18-driven IFN-γ secretion to *T. gondii* depends on parasite invasion but is independent of secreted GRA proteins

Next, we assessed if rapid IFN-γ secretion in response to *T. gondii* required active parasite invasion or could be induced by soluble factors. To this end, naïve B6 mice were injected with either live, heat-killed or sonicated *T. gondii* ME49 tachyzoites. Only inoculation with live parasites induced IFN-γ secretion and increased serum IL-18 levels (**Fig 5A, B**). To exclude the possibility that heat inactivation and sonication destroyed soluble factors that could potentially drive this response, we also injected naïve B6 mice with HFF cell debris, which had been re-suspended in the *T. gondii* ME49 culture supernatant. This treatment also failed to induce IFN-γ and IL-18 secretion (**Fig 5A, B**). These results indicated that active parasite invasion is required to initiate an IFN-γ response, suggesting that *T. gondii* virulence factors may play a critical role. Evidence from studies that have investigated the mechanistic framework of how intracellular bacterial pathogens activate inflammasomes *in vivo*, suggests that secreted effector molecules and/or distinct structural proteins are critically required [34]. Apicomplexan parasites also secrete effector molecules with distinct host-modulatory properties [35]. In particular, dense granule (GRA) proteins have been shown to play important roles in the maintenance of the parasitophorous vacuole (PCV), for the intracellular lifestyle and to exert host-modulatory functions [36]. We further probed the parasite-derived factors that might drive early, IL-18-dependent IFN-γ secretion by exposing naïve B6 mice to a panel of *T. gondii* strains to test if GRA proteins are required for IL-18-driven IFN-γ secretion. Hence, we infected mice with a mutant strain of *T. gondii* ME49 that lacks ASP5, a critical requirement for secretion of GRA proteins [37], as well as strains lacking GRA20 or GRA23, two proteins that contain the PEXEL motif required for PCV exit. No significant difference in the levels of serum IL-18 and NK cell-produced IFN-γ was observed 24 and 48 hours after inoculation with *T. gondii* ME49 ASP5-deficient tachyzoites compared with inoculation of a wildtype *T. gondii* ME49 (**Fig 5C, D**), suggesting that ASP5-driven GRA export is dispensable for inflammasome activation. Similarly, inoculation with GRA20-deficient or GRA23-deficient parasites did not significantly reduce IFN-γ secretion in the absence of IL-12 (**S2A Fig**). We also tested another Type II *T. gondii* strain, DEG (*T. gondii* DEG), which had been implicated in reduced IL-1β secretion following *in vitro* infection of macrophages [24] but, similar to inoculation with *T. gondii* ME49 ASP5-deficient parasites, inoculation with *T. gondii* DEG did not lead to reduced levels of serum IL-18 and NK cell-produced IFN-γ in this model (**Fig 5C, D**). At 48 hours after tachyzoite inoculation, the levels of serum IL-18 were even significantly higher compared with inoculation of *T. gondii* ME49 (**Fig 5D**). These data indicate that ASP5-dependent secretion of GRA proteins does not affect IL-18-driven IFN-γ secretion and highlights the diverging mechanisms that underlie *in vitro* IL-1β and *in vivo* IL-18 secretion in response to *T. gondii*.

**Figure 5:**
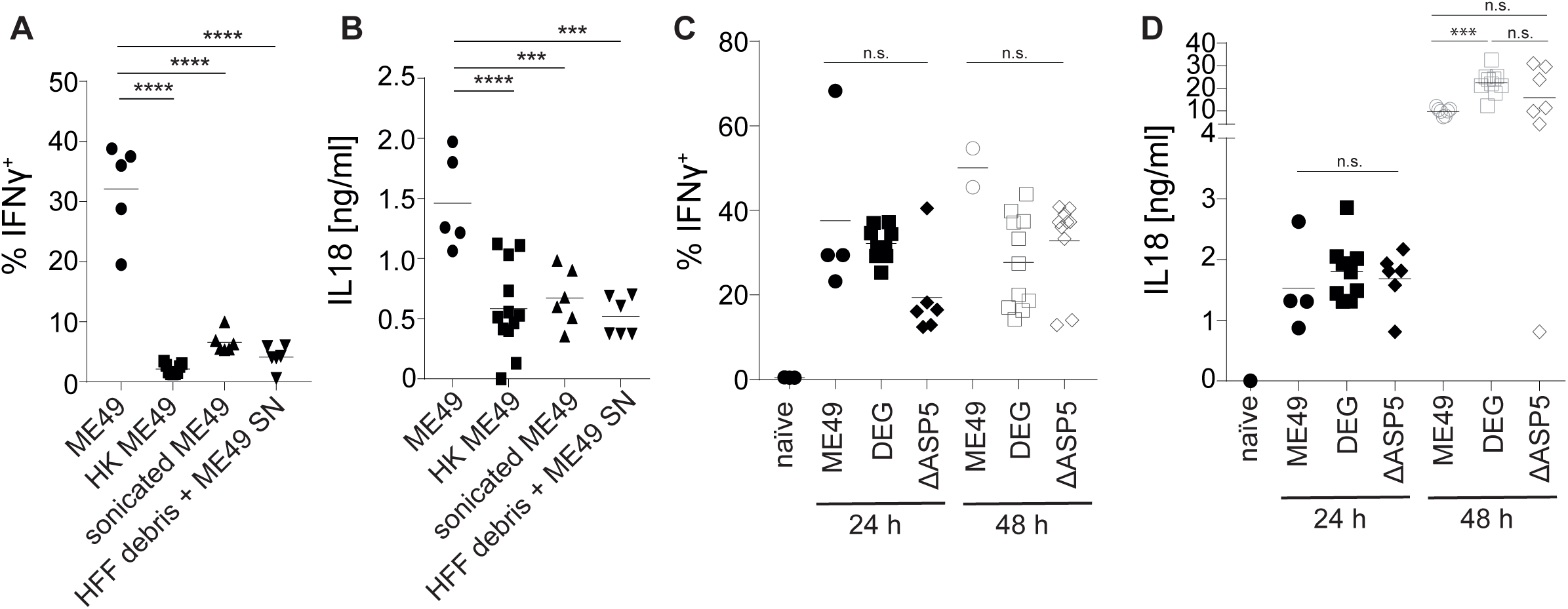
IL-18 driven IFN-γ secretion to *T. gondii* depends on parasite invasion but is independent of secreted GRA proteins. Percent of IFN-γ^+^ cells amongst total viable splenic CD3^−^NKp46^+^ cells (A, C) and serum IL-18 levels (B, D) in naïve mice 24 and 48 hours after i.v. injection of live 10^7^ *T. gondii* ME49 (A-D), DEG (C, D) or ME49ΔASP5 tachyzoites (C, D), heat-killed (A, B) or sonicated ME49 tachyzoites (A, B), or HFF debris with culture supernatant (A, B). Results are presented as individual data points of 4-15 mice per group from at least two pooled independent experiments. Statistical analyses: One-way ANOVA per time-point followed by Dunnett’s multiple comparison test; significant differences are indicated by asterisks: *** p<0.001; n.s. not significant. See also S2 Figure.

### IL2C treatment expands IL-18-responsive IFN-γ-secreting cell subsets

Collectively, the results presented so-far raise the prospect that, if the ability of non-CD4 cells to invoke inflammasome-dependent, IL18-driven production of IFN-γ can be enhanced, it may be possible to control acute toxoplasmosis in AIDS. Hence, we investigated if targeted expansion of non-CD4 cells with IL2C treatment can achieve this. First, naïve mice were treated i.p. with IL2C complex on four consecutive days (**Fig 6A**) and, 24 hours after the last IL2C injection, immune cell expansion was assessed by flow cytometry relative to untreated animals. As reported previously [38], IL2C treatment led to a significant expansion of memory CD8^+^ T cells, NK cells and DN T cells in spleen and MLN (**Fig 6B, C**) and to a minor increase in the Peyer’s Patches (PP) (**Fig 6D**).To further assess if IL2C-expanded and non-expanded CD8^+^ T cells, DN T cells and NK cells responded similarly to *T. gondii* infection, IL2C-treated and untreated mice were infected with 10^7^ ME49 tachyzoites for 24 hours (**Fig 6A**). The percentage of CD8^+^ T cells, DN T cells and NK cells producing IFN-γ was almost indistinguishable between IL2C-treated and untreated mice (**Fig 6E**; data for CD8^+^ T cells and DN T cells not shown). The number of IFN-γ^+^ NK cells (**Fig 6F**), IFN-γ^+^ CD8^+^ T cells and IFN-γ^+^ DN T cells (**S3A Fig**) increased 3-30 fold following IL2C treatment. Similarly, IL2C pretreatment significantly increased systemic IFN-γ levels in the serum after i.v. infection (**Fig 6G)**, but as expected did not lead to a significant change in the levels of upstream serum IL-18 **(S3b Fig**). We also assessed the expression of IL18R and IL12R on the surface of IFN-γ^+^ and IFN-γ^-^ cells. IFN-γ^+^ NK cells (data for CD8^+^ T cells and DN T cells not shown) expressed significant higher levels of IL18R and IL12R compared to IFN-γ^-^ NK cells (**Fig 6H, I**). Taken together, these results show that IL2C-expanded cells respond identically to non-expanded cells and that IL2C treatment numerically expands IFN-γ producing cells that maintain a higher IL18R level expression compared to IFN-γ^-^ cells.

**Figure 6:**
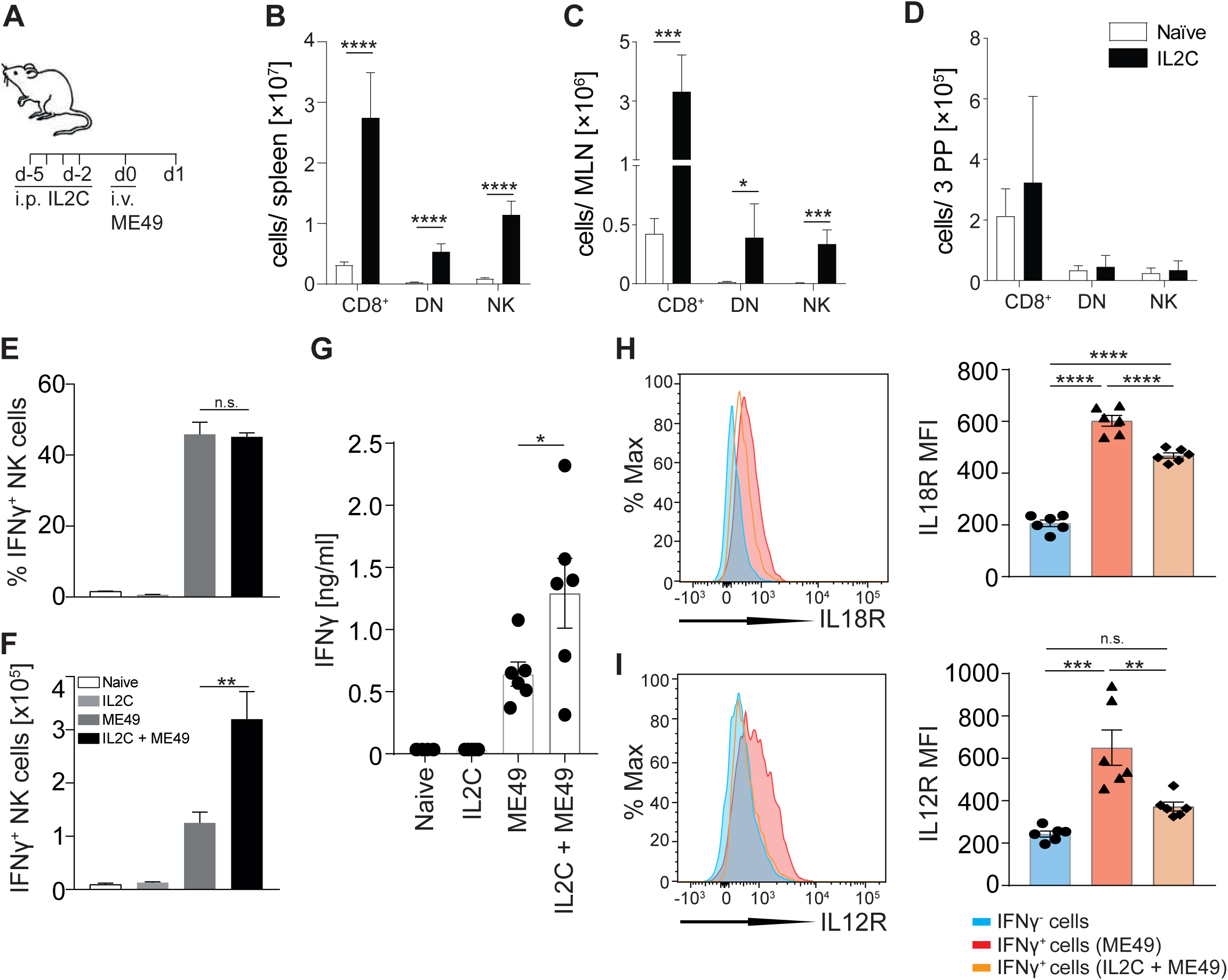
IL2C treatment expands IL-18-responsive IFN-γ-secreting cell subsets. (A-D) Naïve B6 mice were treated i.p. with IL2C on four consecutive days. One day after the last administration, mice were euthanased and numbers of CD3^+^CD8^+^, CD3^+^CD4^−^CD8^−^ (DN) and CD3^-^NKp46+ cells in spleen (B), MLN (C) and PP (D) were assessed by FACS. (E, F) Naïve B6 mice were treated i.p. with IL2C on four consecutive days. Two days after the last IL2C treatment mice were injected i.v. with 10^7^ *T. gondii* ME49 tachyzoites and proportions (E) and total numbers (F) of viable splenic CD3^−^NKp46^+^ IFN-γ^+^ cells were enumerated 24 hours later. (G) IFN-γ serum concentrations 24 hours after mice were injected i.v. with 10^7^ *T. gondii* ME49 tachyzoites. (H, I) Expression of IL18R (H) and IL12R (I) on IFN-γ^-^ (blue histogram) and IFN-γ^+^ CD3^−^NKp46^+^ cells after i.v. infection with 10^7^ *T. gondii* ME49 tachyzoites with (orange histogram) or without (red histogram) IL2C treatment. Results are presented as pooled data means ± SEM with individual data points (G-I) from at least two pooled independent experiments with 5-6 mice per group (B-I) and as representative histograms and individual data points of mean fluorescent intensity (H, I). Statistical analyses: One-way ANOVA followed by Tuckey’s multiple comparison test (A-D); significant differences are indicated by asterisks: * p<0.05; ** p<0.01; *** p<0.001; **** p<0.0001; n.s. not significant. See also S3 Figure.

### IL2C pre-treatment protects mice from acute lethal toxoplasmosis independently of TReg expansion and parasite burden

To definitively assess if IL2C-mediated expansion of IL-18-responsive IFN-γ-secreting non-CD4 cell subsets can prevent lethal toxoplasmosis in mice, we used the well-established oral inoculation model with *T. gondii* ME49 bradyzoite-containing brain cysts. As above, naïve B6 mice were treated i.p. with IL2C for four consecutive days (**Fig 7A**). IL2C treatment was accompanied by a weight loss from which mice recovered within a few days (data not shown). Forty-eight hours after the last IL2C treatment, mice were inoculated orally with 10 or 40 *T. gondii* ME49 cysts and were assessed for weight loss and survival over 60 days. All mice that had been inoculated with 40 cysts and 87% of mice that had been inoculated with 10 cysts, but had not received IL2C injections, succumbed within 14 days after inoculation (**Fig 7B, C**). In contrast, IL2C pre-treatment extended survival in mice that had been inoculated with 40 cysts up to 36 days, and approximately 40% of mice that had been inoculated with 10 cysts survived until day 60 (**Fig 7B, C**).

**Figure 7:**
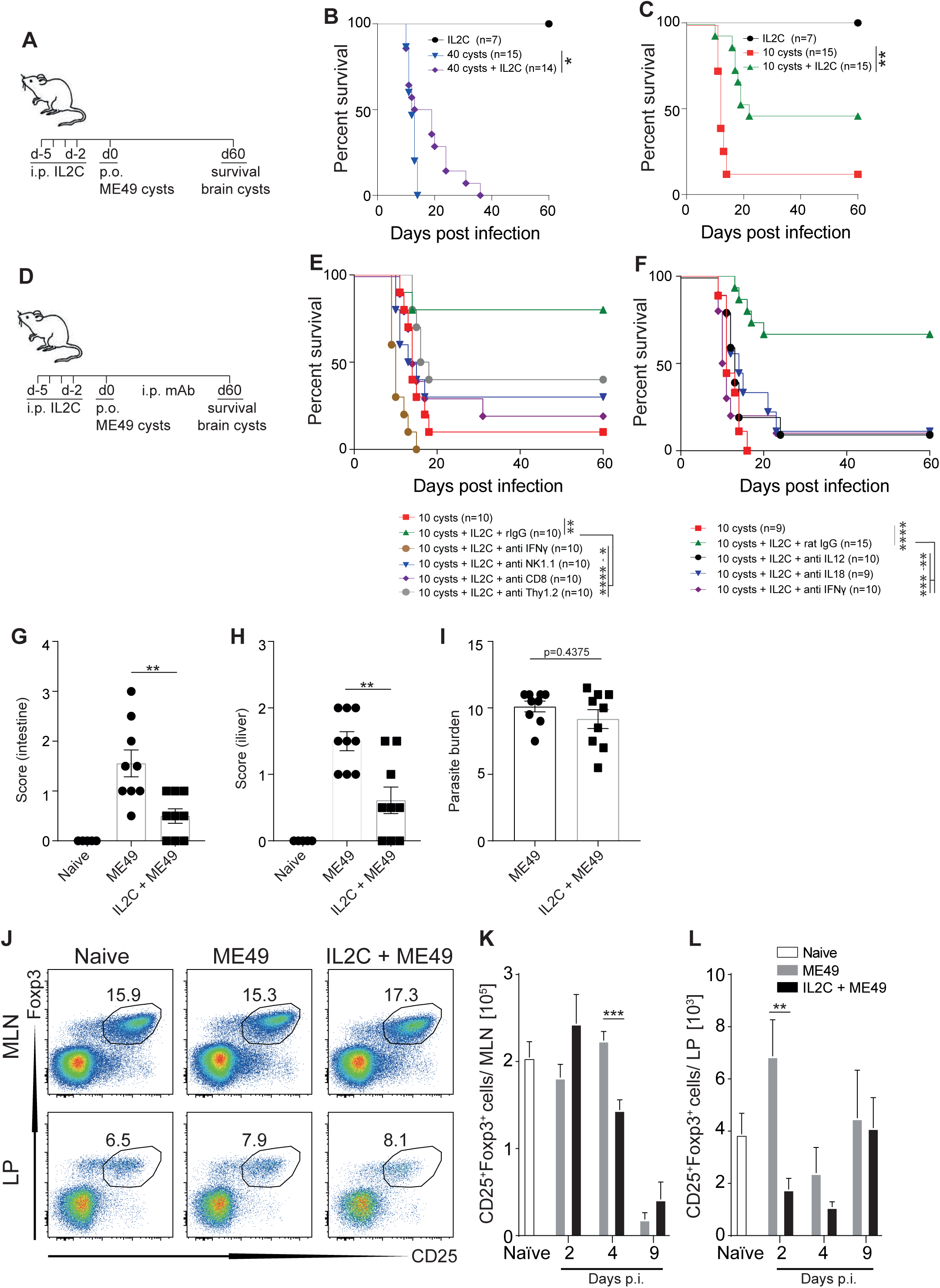
IL2C pre-treatment protects mice from acute, lethal toxoplasmosis independently of TReg expansion and parasite burden. (A-C) Naïve B6 mice were treated i.p. with IL2C on four consecutive days or left untreated. Two days after the last IL2C treatment, mice were inoculated orally with 10 or 40 *T. gondii* ME49 brain cysts and survival was assessed over time. (D-F) Naïve B6 mice were treated i.p. with IL2C on four consecutive days. Two days after the last IL2C treatment, mice were inoculated orally with 10 *T. gondii* ME49 brain cysts. IL2C-treated animals received weekly i.p. injections with mAb against CD8, NK1.1, Thy1.2, IFN-γ or control rIgG (E) or against IL-12, IL-18, IFN-γ (F). Survival was assessed over time (B, C, E, F). Gross pathology of the intestine (G) and liver (H) was assessed 9 days after infection with 10 *T. gondii* ME49 brain cysts. Parasite burden was assessed using splenocytes (I). CD3^+^CD4^+^CD25^+^Foxp3^+^ regulatory T cells were enumerated in MLN and LP at 2, 4 and 9 days after infection. Representative FACS plots from day 2 after infection (J) and mean T_Reg_ numbers ± SEM in MLN (K) and LP (L) are shown. Results are presented as individual data points (G-I), pooled data means (B, C, E, F, K, L) or representative FACS plots (J) from two to three pooled independent experiments with 5-15 mice per group. Statistical analyses: One-way ANOVA followed by Dunnett’s multiple comparison test (G-I, K, L) or Log-rank (Mantel-Cox) test (B, C, E, F); significant differences are indicated by asterisks: * p<0.05; ** p<0.01; *** p<0.001; **** p<0.0001.

Importantly, depletion of NK cells, CD8^+^ T cells, Thy1.2^+^ cells (expressed on all T cells and immature NK cells [39]) or IFN-γ from mice that had been treated with IL2C for four days and had been inoculated with 10 *T. gondii* ME49 cysts with neutralizing antibodies reversed IL2C-mediated increase in survival (**Fig 7D, E**), indicating that IL2C-mediated cell expansion directly correlated with increased survival. Similarly, neutralization of IL-18, IL-12 or IFN-γ, reversed the protective phenotype (**Fig 7D, F**). All mice that were not treated with IL2C succumbed to the infection by day 16, with a median survival of 11 days (**Fig 7F**). Whilst 67% of IL2C-treated mice that received control rat IgG survived until day 60, the median survival for mice treated with anti-IFN-γ was 10.5 days, 13 days for mice treated with anti-IL-12 and 14 days for mice treated with anti-IL-18 (**Fig 7F**). All mice that survived until day 60 were assessed for *T. gondii* brain cysts. Mice contained 100 - 200 cysts per brain (data not shown), indicating that all mice were infected and that survival was not due to a failure of the infection to establish. Taken together, these results further substantiate the proposal that IL2C pre-treatment protects mice from lethal toxoplasmosis *via* IL-12-and IL-18-driven IFN-γ secretion.

To assess if IL2C pre-treatment also impacts on measurable disease parameters other than survival, we also analyzed parasite burden and immunopathology at 2, 4 and 9 days following oral cyst infection. Due to the low infectious dose of 10 cysts, only minimal changes in immunopathology were observed at 2 and 4 days after infection in all groups (data not shown). At 9 days after infection, IL2C pre-treated mice displayed significantly reduced gross pathology of gut and liver (**Fig 7G, H)** in the absence of any effect on parasite burden (**Fig 7I**). TReg numbers in MLN and lamina propria (LP) were not increased after IL2C injections (**Fig 7J, K, L**) suggesting a role for IL2C pre-treatment independent of the previously reported T_Reg_ expansion with JES6-1A12-containing IL2C [40, 41]. Collectively, these results demonstrate a protective role of IL2C pre-treatment in acute lethal murine toxoplasmosis.

## DISCUSSION

Non-CD4 cells, such as CD8^+^ T cells, DN T cells and NK cells, have been implicated in early control of severe infections with intracellular pathogens, including *T. gondii, M. tuberculosis* and *Salmonella* [2, 29]. Our study provides a comprehensive mechanistic framework for how *T. gondii* activates IFN-γ secretion by protective CD8^+^ T cells, DN T cells and NK cells. In particular, we demonstrate that IL-18-driven IFN-γ secretion *in vivo* requires the activation of at least three different inflammasomes; because only *Caspase1/11*^*-/-*^ mice but not *Nlrp1*^*-/-*^, *Nlrp3*^*-/-*^ and *Nlrp1*^*±/-*^*Nlrp3*^*±/-*^ mice were devoid of circulating IL-18 after *T. gondii* infection, a third sensor must exist in addition to NLRP1 and NLRP3 [24, 42]. Furthermore, we show that inflammasome activation occurred in CD8α^+^ DCs, inflammatory monocytes and neutrophils, cell types that have also been implicated in IL-12 secretion in response to *T. gondii* [2]. These results imply a high level of redundancy in the cell type that senses *T. gondii* infection as well as in the host inflammasome signaling pathway. This is in contrast to the often very specific recognition of viral and bacterial infections by one particular inflammasome in a distinct cell type [28, 29, 43–46]. It is likely that this divergence highlights the evolutionary complexity of parasites and suggests that more highly evolved organisms have developed a more complex inflammasome-dependent interplay with their hosts. In line with this hypothesis, it was shown recently *in vitro* that *T. gondii* also activates the NLRC4 and AIM2 inflammasomes in human fetal small epithelial cells [47], as well as the expression of NLRP6, NLRP8 and NLRP13 in THP-1 macrophages [48]. Due to the diverse expression of different internalization receptors and the abundance of inflammasome components, various myeloid cells seem to be endowed with unique abilities to interact with *T. gondii.* In this context the characterization of the myeloid cell populations which produce IL-18 may foster innovative strategies for *T. gondii* interventions.

*Toxoplasma gondii* appears to activate both NLRP1 and NLRP3 [24], yet the specificity of this activation remains elusive. While the activation of NLRP3 in response to *T. gondii* appears to be influenced by P_2_X_7_ receptor-dependent potassium efflux and the induction of reactive oxygen species [47, 49–51], the exact mechanisms of how *T. gondii* activates multiple inflammasomes remain enigmatic. In this context it is also interesting to note that *in vitro* infection of mouse macrophages and human monocytes with *T. gondii* only leads to the secretion of IL-1β, but not IL-18 [24, 52]. In contrast, *in vivo* infection in mice leads to significant secretion of IL-18 but not IL-1β [24]. It has even been suggested that *in vitro* infection of human neutrophils leads to evasion of NLRP3 activation and IL-1β secretion [53]. Furthermore, *in vitro* activation of inflammasomes differs between *T. gondii* strains, and is predominantly induced by Type II parasites [24]. These findings suggest that *T. gondii* has evolved sophisticated diverging effector mechanisms to manipulate inflammasome biology in different host cell subsets, and suggest that secreted effector molecules and/or distinct structural proteins may underlie inflammasome activation. It is, therefore, interesting that *Nlrp1*^±/-^*Nlrp3*^±/-^ mice did not show reduced IL-18 secretion after infection with *T. gondii*. It is important to note that in mice the *Nlrp1* locus is on the same chromosome as the *Nlrp3* gene, meaning that the generation of rare double knockout offspring relies on recombination rather than inheritance. It will therefore be important to further investigate the role of *Nlrp*1 and 3 with alternative methods, such as CRISPR/Cas9 and/or chemical inhibition.

Our study has ruled out ASP5-dependent GRA proteins [54], the most abundant family of *T. gondii*-derived effector molecules [35], as the primary activator of inflammasomes. GRA molecules influence several host cell pathways [55] and are required for the transport of small molecules across the parasitophorous vacuole [56]. These results do not exclude GRA proteins that don’t depend on ASP5 for export, and further studies will have to investigate the role of ASP5-independent GRA proteins as well as rhoptry proteins and other surface structures in driving this process. In particular, the recently described MYR1 protein export system [57–59] may be valuable in answering if secreted effector molecules are at all required to initiate inflammasome activation.

It is tempting to speculate that the overall purpose of activating multiple inflammasomes in multiple cell types is to drive an inflammatory host response that mediates the progression of *T. gondii* into the chronic cyst phase, while at the same time preventing activation of parasite-killing mechanisms. *Toxoplasma* can invade and replicate in virtually all nucleated cell types of warm-blooded animals. From an evolutionary perspective, it is not surprising that the arms race between the host and the parasite has led to the evolution of numerous strategies to activate the immune system (from the parasite’s perspective) and to sense the invasion (from the host’s perspective). The fundamental differences between the habitats and the composition of the immune system of susceptible warm-blooded host species may require *T. gondii* to activate as many different inflammasome sensors as possible. It is well established that *T. gondii* requires a pro-inflammatory, IFN-γ-dominated immune response to form cysts [7]. Because transmission is critical for the parasite’s survival and completion of the life cycle, it is maladaptive for *T. gondii* to kill its host. This may explain why IFN-γ neutralization is fatal, because IFN-γ deficiency favors tachyzoite replication and prevents cyst formation. Furthermore, these findings may also explain why *T. gondii* cysts reactivate after HIV co-infection in humans; HIV destroys CD4^+^ T cells, a prime IFN-γ producer. Hence, we reasoned that a viable adjunct therapy in *T. gondii/* HIV co-infection might be achieved by boosting IFN-γ-producing CD8^+^ T cells, DN T cells and NK cells to prevent acute toxoplasmosis.

The development, maturation and maintenance of IL-18 responsive NK cells, CD8^+^ memory T cells and DN T cells relies on IL-2 and IL-15 [60–65]. While the role of IL-15 in immunity to *T. gondii* remains controversial [66, 67], IL-2 deficient mice are highly susceptible to *T. gondii* infection [68], and administration of recombinant IL-2 enhances survival of *Toxoplasma*-infected mice [69, 70]. The activity of these cytokines is mediated through trans-presentation, a mechanism by which the cytokine is presented to the cytokine receptor complex beta and common γ chains in the context of cell-bound high-affinity alpha (α) chains of the cytokine receptor [71, 72]. Consequently, complexing IL-2 with anti-IL-2 (IL2C) or IL-15 with IL-15RαFc (IL15C) significantly enhances the biological activity of these cytokines *in vivo* [31, 73]. Importantly, the binding site of the anti-IL2 clone used in the IL2C determines whether a preferential expansion of regulatory T cells (T_Reg_; anti-IL-2 clone JES6-1A12) or CD8+ T cell, NK cells and DN T cells occurs (anti-IL-2 clone S4B6) [31, 74].

Using JES6-1A12-containing IL2C, Akbar *et al*. [41] showed that selective expansion of T_Reg_ cells in Type I *T. gondii* RH-infected animals improved control of the parasite. It was also demonstrated that T_Reg_ expansion with JES6-1A12-containing IL2C can overcome the competition for bioavailable IL-2 by regulatory and effector T cells, leading to reduced immunopathology and morbidity during acute Type II *T. gondii* ME49 infection [40]. These studies are in line with other reports showing a collapse of T_Reg_ cells during acute *T. gondii* infection due to IL-2 starvation and an overall protective role of T_Regs_ in acute *T. gondii*-mediated immunopathogenesis [75–78]. In contrast to JES6-1A12-containing IL2C, S4B6-containing IL2C has been shown to boost NK cell and memory CD8^+^ T cell numbers in mice and to enhance their cytolytic capacity against viral infections, malaria [79] and cancer cells [71, 80–82]. Short-term exposure of naïve mice to IL2C containing S4B6 has also been shown to enhance resistance and immunity against *Listeria monocytogenes* infection [83]. Our study is the first to show a protective effect of S4B6-containing IL2C pre-treatment in toxoplasmosis and our results suggest that IL2C pre-treatment can protect mice from lethal toxoplasmosis via distinct mechanisms, depending on the IL-2 mAb clone used to prepare the cytokine complex. Thus, JES6-1A12-contaning IL2C seems to compensate for the limited bioavailability of IL-2 for Treg survival during acute *T. gondii* infection, leading to reduced immunopathology, whereas S4B6-containing IL2C, whilst also reducing pathology without affecting parasite burden, does so in a Treg-independent manner. Thus, S4B6-containing IL2C seems to favor survival and expansion of IL-18-driven IFN-γ secretion, possibly driving parasites towards stage conversion and cyst formation. It is, hence, tempting to speculate that both types of complex could have a synergistic effect if applied together.

Cytokine complex-mediated immunotherapy has not only attracted attention in models of infectious diseases but also in the cancer field [84]. IL2C treatment reduces viral load in a mouse model of gamma-herpesvirus infection [85] and impacts positively on mouse melanoma [86] and BCL1 leukemia [87]. More recently, IL2C treatment has also been tested successfully in cancer models in combination with immune checkpoint blockade [88]. IL-15/IL-15Rα-Fc complexes (IL15C) have also been shown to expand CD8^+^ T cell, DN T cell and NK cell populations, and to protect mice against cerebral malaria via the induction of IL-10-producing NK cells [79]. Whether IL15C would also be protective in our model of lethal toxoplasmosis remains to be investigated. Taken together, these results suggest that cytokine complex treatment may be a more broadly applicable adjunct therapy in infectious diseases, but also highlight that the protective mechanisms may differ between different pathogens and cytokine complex types used. To our knowledge, no data are available yet on any clinical use of IL2C and IL15C in humans. It will be important to consider the hyper-inflammatory response that can be attributed to IL2C and IL15C treatment and, hence, careful consideration should be taken before using cytokine complexes clinically in the context of toxoplasmosis.

In summary, here we delineate the mechanistic framework of how IFN-γ is produced by non-CD4 cell types *in vivo* in response to *T. gondii*, including a crucial role for parasite viability, active invasion and inflammasome-dependent IL-18 secretion. We demonstrate that *in vivo* inflammasome activation in response to *T. gondii* occurs in multiple myeloid cell types and involves at least three different redundant inflammasomes. Additionally, our study excludes *T. gondii*-derived, ASP5-dependent, dense granule proteins as the main activators of inflammasomes *in vivo*. The observation that both IL-12 and IL-18 neutralization reverses the host protective role of CD8^+^ T cells, DN T cells and NK cell-produced IFN-γ during *T. gondii* infection highlights the redundancy and functional interchangeability of both cytokines during *T. gondii* infection. This combination of observations led us to the hypothesis that enhancement of inflammasome-dependent, IL18-driven production of IFN-γ by non-CD4 cells may be a route to control acute toxoplasmosis in AIDS. Consequently, we provide compelling evidence for a protective role of IL2C pre-treatment in lethal toxoplasmosis. We demonstrate that IL2C-mediated expansion of CD8^+^ T cells, NK cells and DN T cells protects mice against acute disease and death in an IFN-γ-dependent manner. Hence, we conclude that inducing immune responses that lead to the expansion of CD8^+^ T cells, DN T cells and NK cells in combination with inflammasome-dependent, IL-18-driven IFN-γ secretion could be a crucial feature of improved toxoplasmosis intervention strategies, particularly in the context of HIV co-infection.

## Materials and Methods

### Mice

C57BL/6J and Arc(S) mice were purchased from the Animal Resource Center (Perth, Australia). Knockout mice (*Caspase1/11*^−/−^, *Nlrp1*^*-/-*^, *Nlrp3*^−/−^ and *Il18*^*-/-*^) were bred and maintained at the Australian Institute of Tropical Health and Medicine, James Cook University, Cairns and Townsville, Australia. Double knockout mice (*Nlrp1*^*-/-*^*Nlrp3*^*-/-*^) mice were bred by sequentially crossing *Nlrp1*^*-/-*^ [89] and *Nlrp3*^*-/-*^ mice. Genotyping was performed using the following primer pairs: *Nlrp3*-F 5’-GCTCAGGACATACGTCTGGA-3’; *Nlrp3*-R 5’-TGAGGTCCACATCTTCAAGG-3’; *Nlrp3*-R2 5’-TTGTAGTTGCCGTCGTCGTCCTT-3’; *Nlrp1* WT: *Nalp1a*F 5’-TGGAAGGAAGGCAAGCTTTA-3’; *Nalp1a*R 5’-ACCCAGGGAACTTCACACAG-3’; *Nlrp1* mutant: *Nalp1a*F 5’-TTTAGAGCTTGACGGGGAAA-3’; *Nalp1a*R 5’-GGAAGGACTTCCCACCCTAA-3’. The following mice were used for experiments: *Nlrp1*^*-/-*^*Nlrp3*^*-/-*^, *Nlrp1*^*+/-*^*Nlrp3*^*-/-*^ and *Nlrp1*^*-/-*^*Nlrp3*^*+/-*^. For infection experiments, all mice were sex-and age-matched, and kept in our BSL 2 animal facility under specific pathogen-free (SPF) conditions.

### Parasites

Type II *T. gondii* strains ME49, ME49-RFP, ME49 GRA20-deficient, ME49 GRA23-deficient, ME49 ASP5-deficient and DEG (ATCC, ATC50855) were maintained by continuous passage in human foreskin fibroblasts (HFF; ATCC, ATCSCRC1041) in DMEM supplemented with 10% FCS, penicillin, streptomycin and L-glutamine at 37°C and 5% CO_2_. Parasites were harvested from recently lysed cell monolayers, passed through a 26G needle and a 3 µm TSTP Isopore^TM^ membrane filter and concentrated by centrifugation at 500*g* for 10 minutes. The pellet of tachyzoites was re-suspended in sterile PBS. Parasites were counted using a Neubauer hemocytometer and diluted to the required infectious dose in sterile PBS.

### Generation of *T. gondii* ME49 *Gra20* and *Gra23* knockouts

We employed a CRISPR/Cas9 approach to insert frameshifts within the first 20 nt of the start of the coding sequence of *gra20* and *gra23* in *T. gondii* Me49 with consequential disruption of the final translated proteins. Inverse PCR was used to exchange the sgRNA of UPRT with the sgRNA for GRA20 with Ph-sgRNA_TgGRA20mutF (5’-ATGCATAGCCGGAACTGCGTGTTTTAGAGCTAGAAATAGC-3’) and Ph-genCas9mutR (5’-AACTTGACATCCCCATTTAC-3’) to yield plasmid pCAS9sgGRA20. Similarly, inverse PCR was used to exchange the sgRNA of UPRT with the sgRNA for GRA23 with Ph-sgRNA_TgGRA23mutF (5’-GCAGCGCGTGCGGGAAGCAGGTTTTAGAGCTAGAAATAGC-3’) and Ph-genCas9mutR (5’-AACTTGACATCCCCATTTAC-3’) to yield plasmid pCAS9sgGRA23. Transfection of *T. gondii* Me49 was carried out as described previously [90]. Twenty-four hours post-transfection, transiently transfected GFP^+^ parasites were purified by flow cytometry as previously described [91] and individual GRA20 and GRA23 KO clones were further purified using two rounds of limiting dilution cloning. Sanger sequencing of PCR products was used to confirm disruption of the *gra20* and *gra23* ORFs.

### Infections

To isolate *T. gondii* ME49 bradyzoite containing cysts, the brains of chronically infected Arc(S) mice (injected i.p. with 500 tachyzoites of *T. gondii* ME49 >8 weeks prior) were removed, homogenized in sterile PBS, and subjected to centrifugation in a discontinuous Percoll gradient. Cysts were counted using a Neubauer hemocytometer and diluted in sterile PBS. For experiments, B6 mice were inoculated with 10, 40 or 100 cysts by oral inoculation. For mechanistic studies, B6 mice were injected i.v. in the lateral tail vein with 10^7^ tachyzoites of *T. gondii* ME49, mutant strains on the *T. gondii* ME49 background or the Type II strain *T. gondii* DEG in a volume of 200 µl. For heat inactivation, *T. gondii* ME49 tachyzoites were grown as described above, enumerated, and washed twice with PBS before incubation at 62° C in a water bath for 1 hour. Effective killing was verified by addition of heat-killed parasites to a HFF cell monolayer.

### Isolation of leukocytes

Spleens, mesenteric lymph nodes and Peyer’s Patches were extracted and mechanically disrupted by pushing cells through a 70 µm cell strainer. Subsequently, red-cell depleted, single-cell suspensions were prepared as described elsewhere [39]. Lamina propria cells were isolated from the ileum as published previously with minor modifications [92].

### Scoring of pathology

Gross pathology of ileum and liver was scored visually using a scoring system adapted from Melgar et al. [93]. For the ileum, the consistency of the intestinal contents, the degree of swelling and amount of angiogenesis were assessed. This system is based on an ascending scale of severity, for each parameter, as follows: 0 (no abnormality); 1 (minimal); 2 (moderate); or 3 (severe). For the liver, the colour and appearance of the organ were assessed on an ascending scale of severity from 0 (normal colour and appearance); 1 (blotchy appearance with some areas exhibiting change in colour); 2 (entire organ pale in colour); or 3 (entire organ pale in colour with visible signs of necrosis). Scores for each parameter were added together to give a total score for each animal.

### Parasite burden

Parasite burden was measured in the whole spleen of individual mice using a microtitre dilution method adapted from Buffet et al. [94] It was necessary to determine parasite burden in the spleen rather than the intestine because it was impossible to harvest immune cells for analysis from the intestine and determine parasite burden in the same animal; however, we have demonstrated previously that the parasite burden in the spleen accurately mirrors that in the intestine [50]. Briefly, prior to the experiment, 96 well plates were seeded with HFF cells and allowed to become confluent. One row was allocated per mouse and each mouse was done in duplicate. Spleens were removed and single-cell suspensions were made by passing through a 70-µm cell strainer. Cells were pelleted at 1500*g*, and then resuspended in RPMI 1640 containing 5% FCS at a concentration of 1×10^7^ cells/ml. Two hundred microliters of spleen cell suspension was added to the first well of a 96-well plate and then serially diluted 1/2 across the plate. Plates were incubated at 37°C in 5% CO2 for 7 days before wells were examined for the presence of parasites. A score of parasite burden was allocated based on the last column in which parasites were visible.

### Flow cytometry

To assess expression of surface antigens and IFN-γ secretion, viable, red blood cell-depleted single-cell suspensions were stained with monoclonal antibodies (all from BD Pharmingen) against CD4 (clone GK1.5), CD8α (clone 53-6.7), CD3 (clone 145-2C11), NKp46 (clone 29A1.4), CD44 (clone 1M7), CD90.1 (clone 30-H12), CD11b (clone M1/70), CD11c (clone HL3), MHC-II (clone M5/114), CD11b (clone M1/70), Ly6G (clone 1A8), Ly6C (clone AL-21), CD19 (clone 1D3), F4/80 (clone BM8), or IFN-γ detection antibody (Miltenyi Biotec, Germany). After washing the cells, samples were analyzed using a FACSCantoII or FortessaX20 analyzers (BD Biosciences, CA). Propidium iodide (2 μg/ml) was added to exclude dead cells.

### Assessment of *ex vivo* IFN-γ secretion

*Ex vivo* IFN-γ secretion by distinct lymphocyte subsets was assessed as described previously [29]. Briefly, mice were injected i.v., i.p., or p.o. with different doses of *T. gondii* ME49 cysts or tachyzoites (as described in figure legends). At different time points after injection of parasites (as described in figure legends), organs were removed aseptically, single cell suspensions were prepared and red blood cells were lysed. Cells (10^6^) were stained with the ‘Mouse IFN-γ secretion assay detection kit’ (Miltenyi Biotec, Germany) according to the manufacturer’s instructions and IFN-γ secretion was analyzed by flow cytometry.

### Detection of *in vivo* inflammasome activation by flow cytometry

Detection of apoptosis-associated speck-like protein containing a caspase recruitment domain (ASC) assembly was performed as described previously [95]. Briefly, mice were injected with 10^7^ *T. gondii* ME49-RFP tachyzoites and euthanased 24 hours later. Cells were stained for surface molecules, fixed, permeabilized and stained with rabbit anti-ASC antibody (Santa Cruz Biotechnology) for 45 min at room temperature. Subsequently, a secondary anti-rabbit Alexa488 antibody (Life Technologies) was added for 45 min at room temperature. A FMO control without anti-goat Alexa488 was included.

Detection of active caspase-1 by flow cytometry was performed using the carboxyfluorescein FLICA kit (FAM-YVAD-FMK, Immunochemistry Techniques, Bloomington, MN). B6 mice were injected with 10^7^ *T. gondii* ME49-RFP tachyzoites and 23 hours later FAM-YVAD-FMK (diluted in DMSO and PBS) was injected intravenously. Splenic cells were analyzed by FACS 1 hour later as described above (24 hours after *T. gondii* ME49-RFP injection). Mice that received *T. gondii* ME49-RFP but no FAM-YVAD-FMK were used as FMO control. RFP^+^ cells were analyzed for expression of cell specific surface markers and positivity in green fluorescence.

### Multiplex and ELISA

Blood for serum analysis was taken post mortem from the aorta abdominalis and collected in serum separator tubes (BD), left for 30 minutes at room temperature, followed by centrifugation at 12,000*g* for 3 min. Sera were stored at –20°C until analysis. Measurements were performed using CBA (BD Biosciences, CA) or ELISA (elisakit.com, Australia) according to manufacturers’ instructions. Samples were acquired on a FACSCantoII (BC Biosciences, CA) or a FLUOstar Omega ELISA Reader (BMG Labtech).

### IL-2/anti-IL-2 complex-mediated cell expansion

IL-2/anti-IL-2 complexes (IL2C) were prepared as described previously [38]. Briefly, 1.5 µg of recombinant mouse IL-2 (Peprotech) and 10 µg of anti-IL-2 mAb (clone S4B6, Walter and Eliza Hall Institute [WEHI] antibody facility, Melbourne, Australia) were mixed, incubated at 37°C for 30 min, and administered i.p. in a volume of 200 µl for four consecutive days.

### Antibody-mediated cell depletion and cytokine neutralization

For cytokine neutralization and cell depletion, monoclonal antibodies against IL-12, IL-18, IFN-γ, CD8, NK1.1, Thy1.2 and rat IgG were purchased from the WEHI antibody facility or from BioXCell (NH, USA). A total of 200 µg of anti-IL-18 (clone YIGIF74-1G7; Cat. No.: BE0237), anti-IFN-γ (clone HB170-15), anti-IL12 (clone C17.8), anti-NK1.1 (clone PK136), anti-CD8 (clone 2.43), anti-Thy1.2 (clone 30H12) or control rat IgG were injected i.p. weekly in a volume of 200 µl.

### Statistics

Flow cytometry data were analyzed using FlowJo software (Treestar, CA) and statistical analysis was performed using GraphPad Prism, GraphPad software, San Diego, CA as indicated in individual figure legends. One-way analysis of variance (ANOVA) was followed by Dunnett’s multiple comparison test, and two-tailed Student’s *t* tests were used. A Log-rank (Mantel-Cox) test was used to compare significance for survival experiments. A P value of less than 0.05 was considered significant.

### Ethics Statement

All experiments were approved and conducted according to Australian animal protection law and in accordance with requirements of the Animal Ethics Committee of James Cook University (A2138, A2324). Death was never used as an endpoint.

## Acknowledgments

This work was supported by the National Health and Medical Research Council of Australia (NHMRC) through a CJ Martin Biomedical Early Career Fellowship (APP1052764) and a Career Development Level 1 Fellowship (APP1140709) to A.K. A.K and C.M.M. also acknowledge support through a Capacity Building Grant (15031) provided by the Australian Institute of Tropical Health and Medicine (AITHM) *via* the Queensland Government. We thank Michael E. Grigg (NIH, USA) for generously providing the *T. gondii* ME49-RFP, and Seth Masters (WEHI, Melbourne, Australia) for providing *Nlrp1*^*-/-*^ and *Nlrp3*^*-/-*^ mice. We are grateful to Dominique Soldati-Favre (University of Geneva) for her insights into GRA proteins.

## Author contributions

A.K., C.M.M. and N.C.S. conceived the study; A.K., C.M.M, R.A.W., J.A.W., P.R.G. and S.P. performed experiments; P.M.H. and P.R.G. provided reagents and intellectual input. A.K. performed data analysis and wrote the manuscript. N.C.S and C.M.M. commented extensively on the manuscript; all coauthors read and approved the final manuscript.

---

**S1 Figure:**
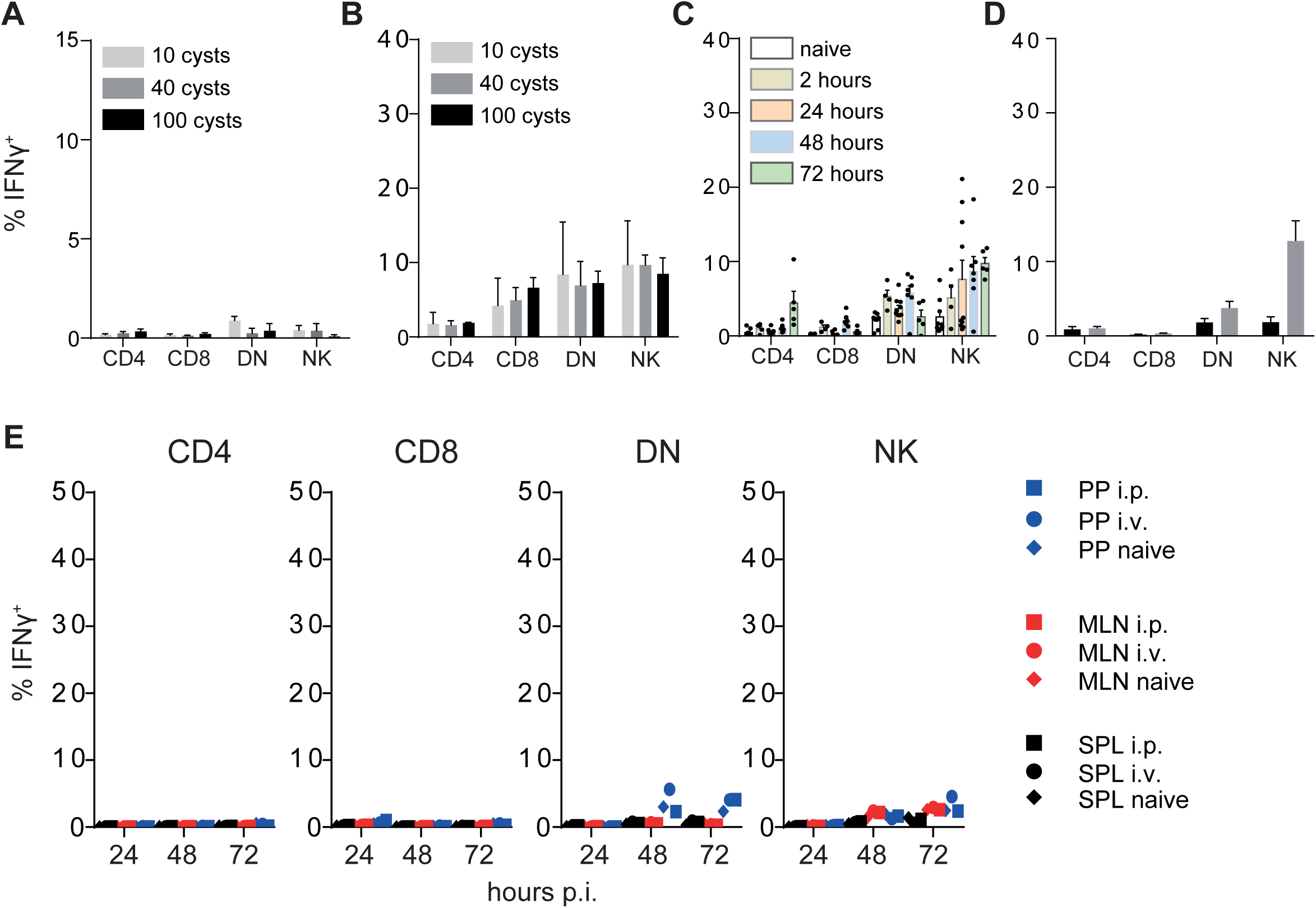
Low dose injection of *T. gondii* ME49 tachyzoites does not induce rapid IFN-γ secretion. Percent of IFN-γ^+^ cells amongst total viable CD3^+^CD4^+^, CD3^+^CD8^+^, CD3^+^CD4^−^ CD8^−^ (DN) T cells and CD3^−^NKp46^+^ cells in three Payers Patches 1 day (A) or 5 days (B) after B6 mice were inoculated orally with 10, 40 or 100 *T. gondii* ME49 brain cysts; 2-72 hours after mice were injected i.v. with 10^7^ *T. gondii* ME49 tachyzoites (C) or 24 hours after mice were infected i.p. with 10^7^ *T. gondii* ME49 tachyzoites (D). (E) Percent of IFN-γ^+^ cells amongst total viable CD3^+^CD4^+^, CD3^+^CD8^+^, CD3^+^CD4^−^CD8^−^ (DN) T cells and CD3^−^NKp46^+^ cells from spleen, mesenteric lymph nodes or three Peyers Patches (PP) at 2-72 hours after B6 mice were injected i.p. or i.v. with 10^5^ *T. gondii* ME49 tachyzoites. Results are presented as individual data points (E) or pooled data means (A-D) from two pooled independent experiments with 3-10 mice per group.

**S2 Figure:**
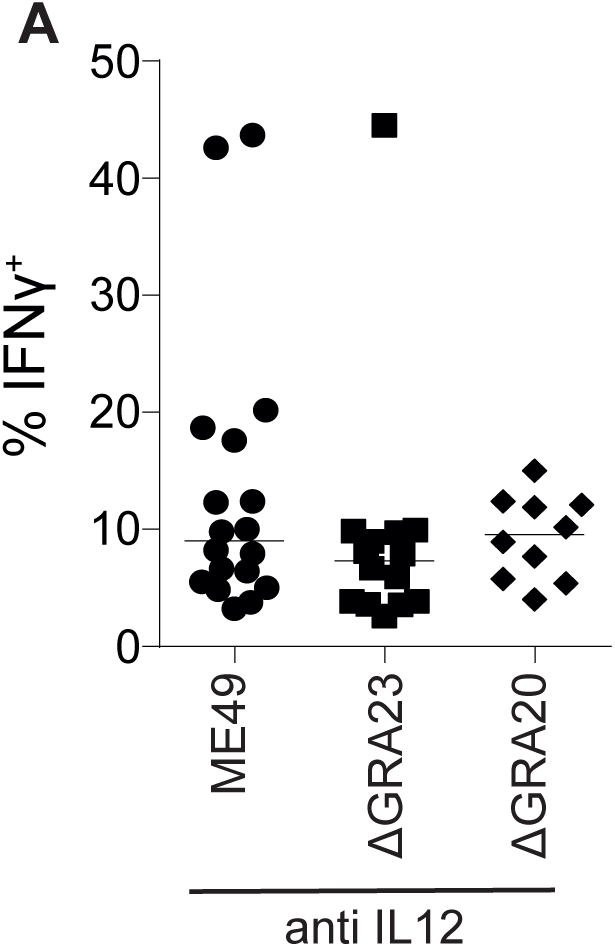
IL-18 driven IFN-γ secretion to *T. gondii* is independent of secreted GRA proteins. Percent of IFN-γ^+^ cells amongst total viable splenic CD3^−^NKp46^+^ cells in naïve mice 24 hours after i.v. injection of 10^7^ *T. gondii* ME49, ME49 GRA20-deficient or ME49 GRA23-deficient tachyzoites. Mice were treated with mAb against IL-12 immediately after injection of *T. gondii*. Results are presented as individual data points of 4-15 mice per group from at least two pooled independent experiments. Statistical analyses: One-way ANOVA followed by Dunnett’s multiple comparison test; not significant.

**S3 Figure:**
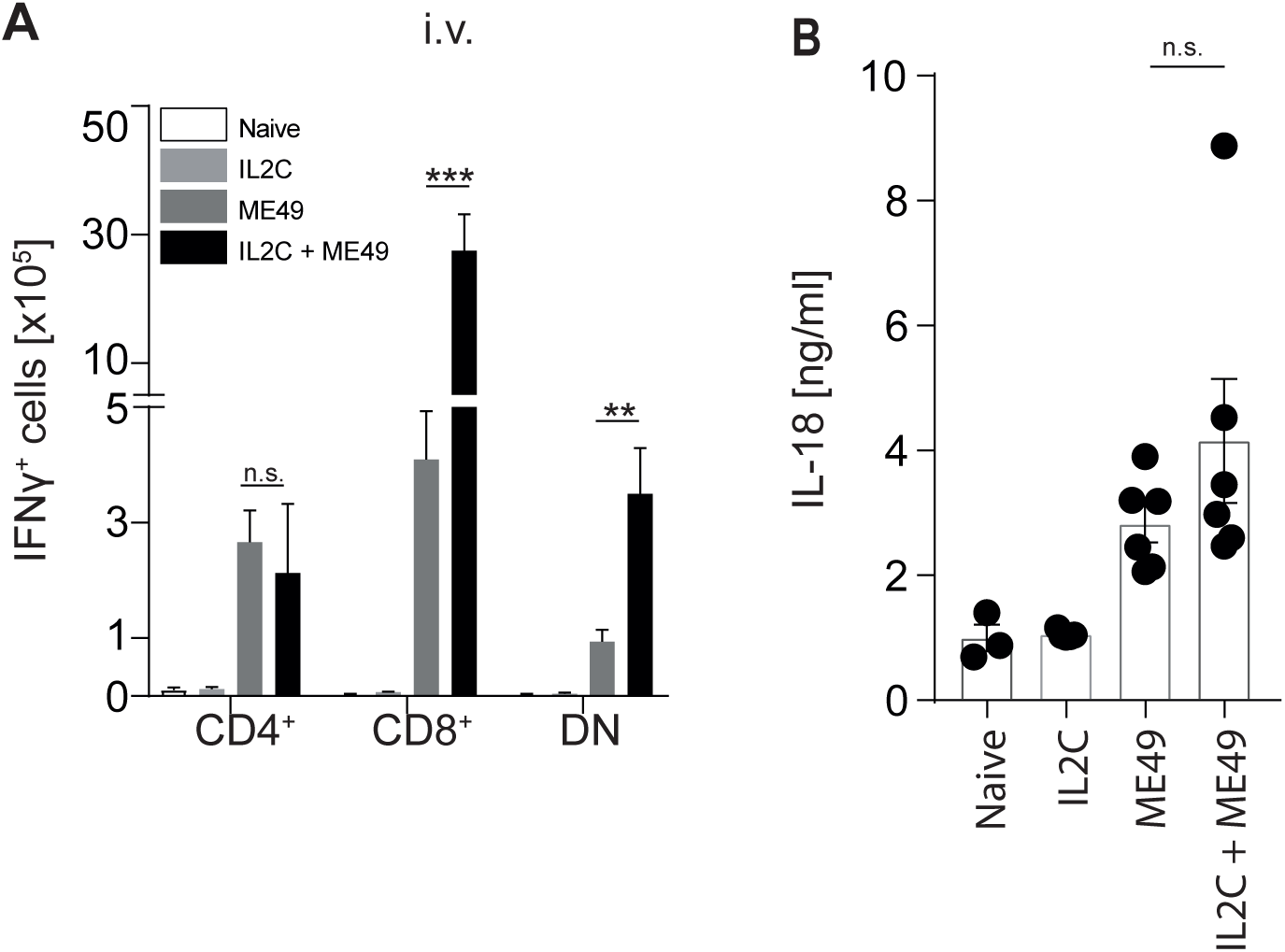
IL2C treatment expands IL-18 responsive IFN-γ-secreting cell subsets but has no impact on upstream IL-18 secretion. (A, B) Naïve B6 mice were treated i.p. with IL2C on four consecutive days. Two days after the last IL2C treatment mice were injected i.v. with 10^7^ *T. gondii* ME49 tachyzoites and viable splenic CD3^+^CD4^+^, CD3^+^CD8^+^, CD3^+^CD4^−^CD8^−^ (DN) T cells IFN-γ-secreting cells (A) were enumerated 24 hours later, and serum IL-18 (B) levels were measured. Statistical analyses: One-way ANOVA followed by Dunnett’s multiple comparison test; significant differences are indicated by asterisks: * p<0.05; ** p<0.01; *** p<0.001; n.s. not significant.

